# Evaluation of PECAM-1 Expression and Microvessel Density in Gastric Adenocarcinoma: a Cross-sectional Study

**DOI:** 10.1101/2025.07.30.667720

**Authors:** Gazi Abdus Sadique, Tanshina Afrin

## Abstract

**Background:** Gastric adenocarcinoma is the fifth most prevalent malignancy globally and ranks sixth in Bangladesh, representing a significant oncological and public health challenge. Gaining in depth knowledge about the tumor microenvironment, particularly the mechanisms of tumor-induced angiogenesis, is essential for the development of more precise and effective targeted therapies. Microvessel density (MVD) serves as a widely recognized measure of angiogenic activity and can be reliably assessed using immunohistochemical staining for PECAM-1 (CD31), a highly specific marker of vascular endothelial cells.

**Methods:** This cross-sectional observational study was conducted in the Department of Pathology, Satkhira Medical College, from April 2024 to March 2025. A total of 50 cases of invasive gastric adenocarcinoma were included. Routine Hematoxylin and Eosin (H&E) staining was performed for Lauren classification. Immunohistochemistry for PECAM-1 was conducted to highlight microvessels.

**Result:** Among 50 cases, intestinal-type adenocarcinoma was more frequent than diffuse type. High MVD was observed in 58.1% of intestinal-type cases and 36.8% of diffuse-type cases. The difference in MVD between intestinal and diffuse types was statistically significant (p < 0.05), suggesting higher angiogenic activity in intestinal-type tumors.

**Conclusion:** This study demonstrates that PECAM-1 positive MVD is significantly higher in the intestinal subtype of gastric adenocarcinoma compared to the diffuse subtype. These findings indicate a more angiogenically active tumor microenvironment in intestinal-type tumors, potentially correlating with greater invasive potential and metastatic behavior. PECAM-1 immunostaining provides a valuable tool for quantifying tumor angiogenesis and may serve as a prognostic marker or a basis for anti-angiogenic therapeutic targeting in gastric cancer management.

## Background

Gastric adenocarcinoma ranks as the fifth most common cancer worldwide and the sixth in Bangladesh, posing a serious public health and oncologic burden. Despite recent advancement in treatment strategies, outcomes remain disappointing, especially in patients with advanced disease [1]. A deeper understanding of the tumor microenvironment, particularly the tumor angiogenesis, is key to identifying more effective, targeted therapies [2]. Angiogenesis is the development of new blood vessels from existing ones. It is a driving force behind tumor growth, invasion, and metastasis [3]. A commonly used indicator of angiogenic activity is microvessel density (MVD), which can be accurately measured through immunohistochemical analysis of PECAM-1 (CD31) which is a marker highly specific to endothelial cells [4].

In this study, we examined PECAM-1 expression and microvessel density across two common histological subtypes of gastric adenocarcinoma. Our goal was to evaluate their role in tumor aggressiveness and investigate their potential utility in guiding therapeutic approaches.

## Materials and Methods

This cross-sectional descriptive study was conducted in the Department of Pathology, Satkhira Medical College, from April 2024 to March 2025. Ethical approval was obtained from the Ethical Review Committee of Satkhira Medical College prior to the initiation of the study. Routine Hematoxylin and Eosin (H&E) staining was performed at the Department of Pathology, Satkhira Medical College. Immunohistochemical analysis for PECAM-1 (CD31) was carried out at the Department of Pathology, Bangabandhu Sheikh Mujib Medical University (BSMMU), Dhaka to assess tumor angiogenesis. A total of 50 histopathologically confirmed cases of gastric adenocarcinoma were included. Tissue samples were obtained through gastrectomy and endoscopic biopsies, and relevant clinical and histopathological data were collected using a pre-tested structured questionnaire.

Weidner’s two dimensional counting method was used for MVD interpretation [5]. CD31 staining showed a single stained endothelial cell or a cluster of endothelial cells as a positive microvessel. Firstly, the regions with the most abundant vasculature in tumor margins and tumor center, namely “hot spots” were selected under low magnification, and then 5 fields were randomly counted under high magnification (200 times), with their mean value as MVD count. Any endothelial cell or group of endothelial cells that were stained brown and adjacent to the vessel could be used as a counting microvessel. Vessels with a lumen larger than 8 red blood cells and a thicker wall were not counted.

## Results and observation

50 histopathologically diagnosed gastric adenocarcinoma cases were taken. Patients (case) with age ranged from 30 to 66 years (mean age was 53.34 ± 8.38), 64% (n=32) patients were male, and 36% (n=18) patients were female. Out of 50 gastric adenocarcinoma cases, 31 (62%) cases were intestinal type, and 19 (38%) cases were diffuse type adenocarcinoma. Mean MVD in the study group was 17.44 ± 9.88, with a median value of 13.50. High MVD was found in 18 (58.1%) cases of intestinal type of gastric adenocarcinoma and in only 07 (36.8%) cases of diffuse type of gastric adenocarcinoma. The MVD count in intestinal type gastric cancer was significantly higher than that in diffuse type gastric cancer (P < 0.001).

**Figure 1.**
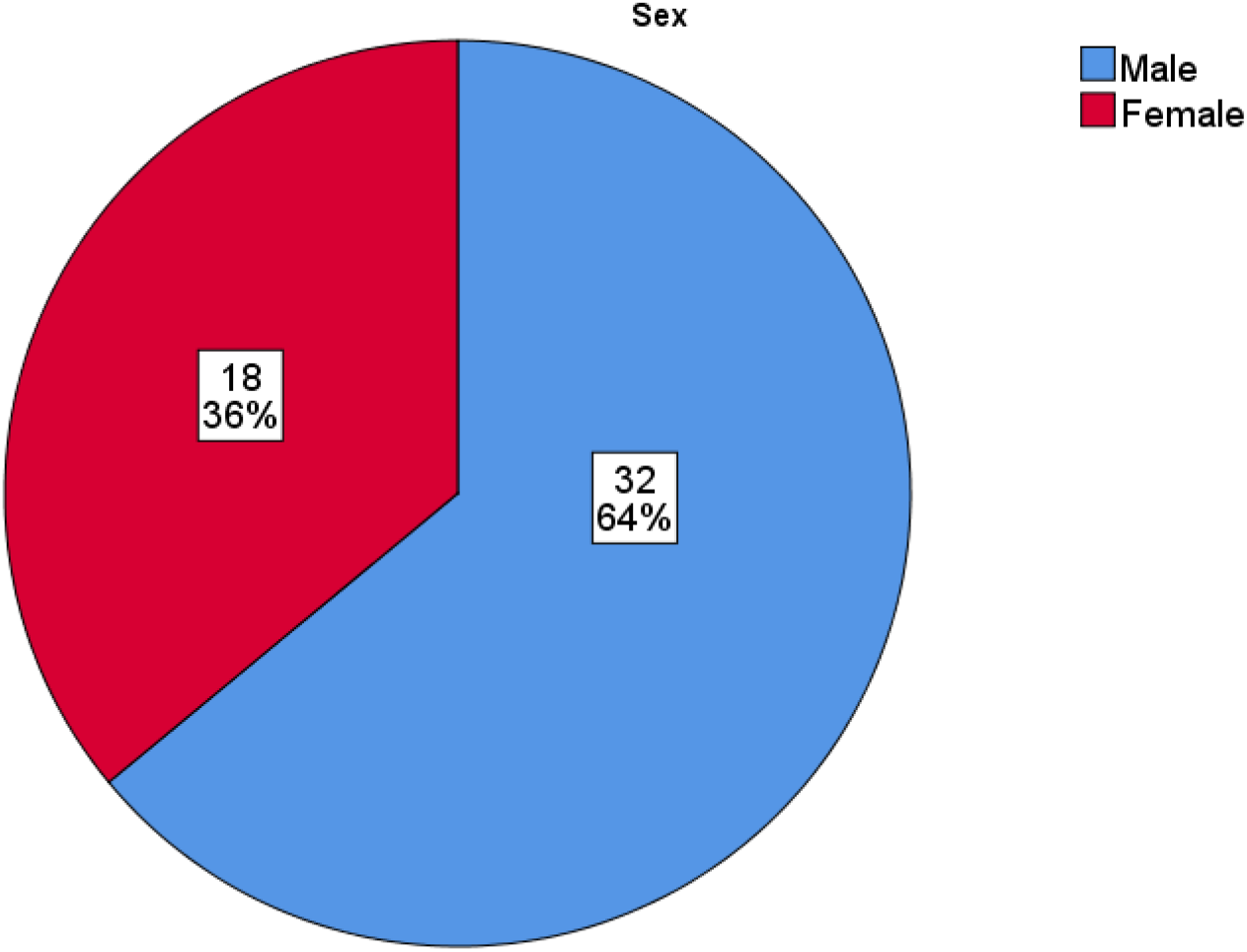
Distribution of cases according to gender (n=50)

**Figure 2.**
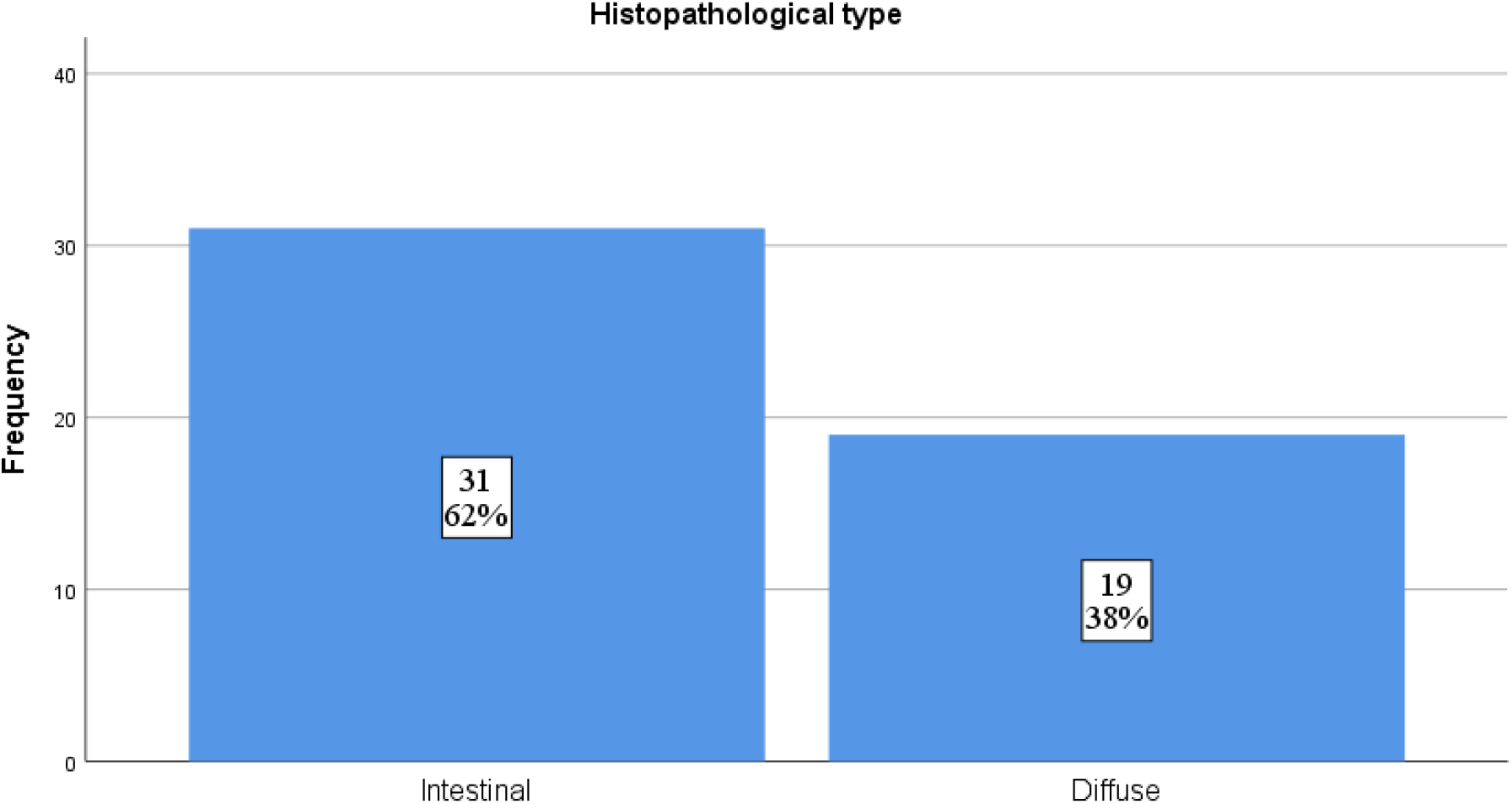
Distribution of tumor type among the study cases (n=50)

**Figure 3.**
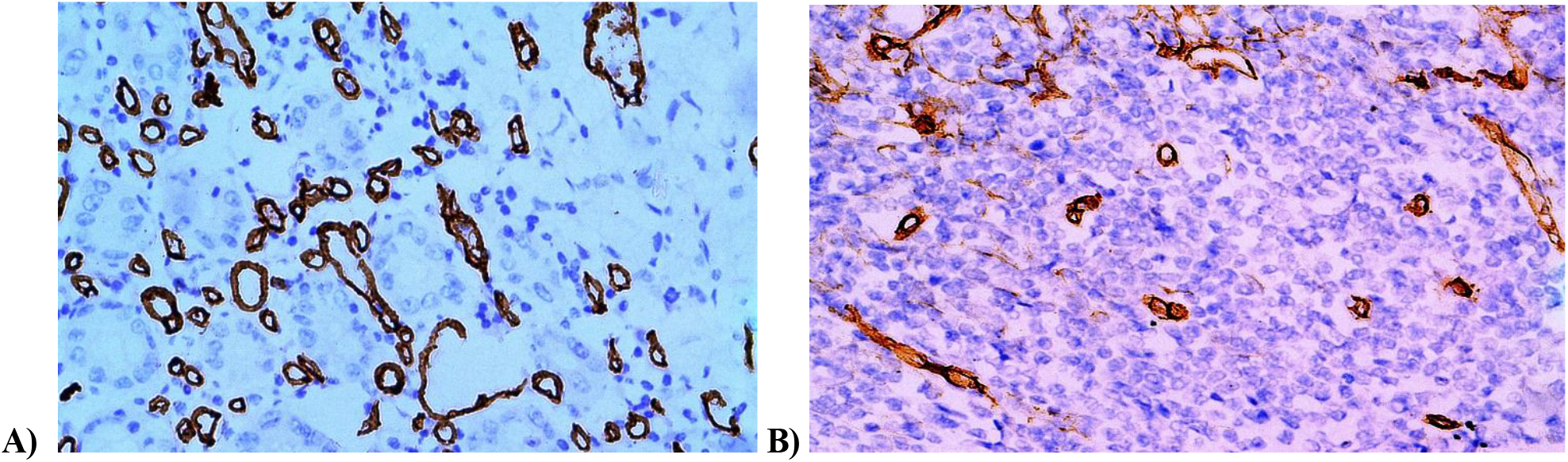
A. Higher MVD counts in intestinal type of gastric adenocarcinoma B. MVD counts were fewer in diffuse type of gastric adenocarcinoma.

**Table 1.**
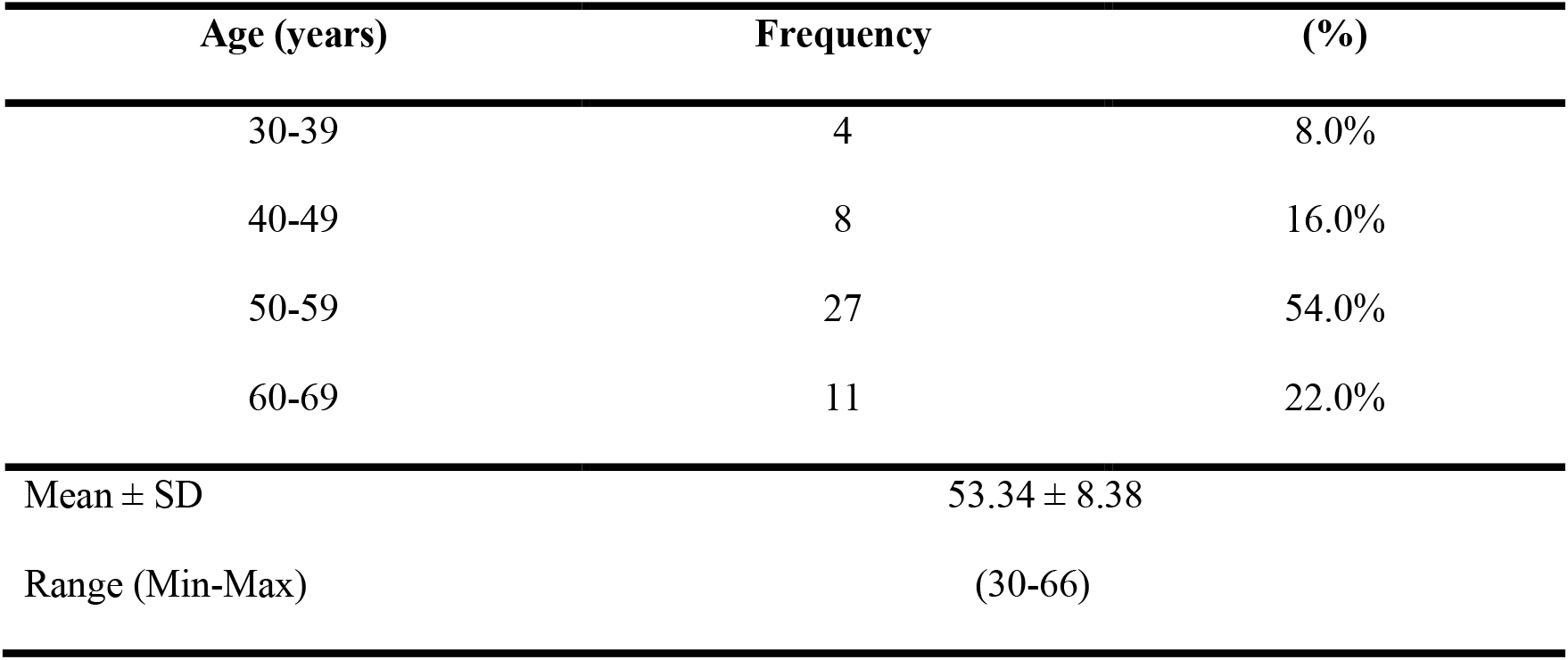
Distribution of the study subjects by their age (n = 50).

| Age (years) | Frequency | (%) |
| --- | --- | --- |
| 30-39 | 4 | 8.0% |
| 40-49 | 8 | 16.0% |
| 50-59 | 27 | 54.0% |
| 60-69 | 11 | 22.0% |
| Mean ± SD | 53.34 ± 8.38 | |
| Range (Min-Max) | (30-66) |  |

**Table 2.** Microvessel density among the study groups (n=50).

| Microvessel density | Frequency | Percent |
| --- | --- | --- |
| Low | 25 | 50.0 |
| High | 25 | 50.0 |
| **Mean ± SD** | **17.44 ± 9.88** |  |
| **Median** | **13.50** |  |
| **Range (Min-Max)** | **(6.00-49.00)** |  |

**Table 3.** Association of MVD with histopathological types of gastric adenocarcinoma (n=50).

| MVD status | Histopathological type |  | p-value |
| --- | --- | --- | --- |
|  | Intestinal | Diffuse |  |
| High | 18 | 7 | **0.049** |
|  | 58.1% | 36.8% |  |
| Low | 13 | 12 |  |
|  | 41.9% | 63.1% |  |

\*Data were analyzed using **Pearson Chi-Square Test**.

## Discussion

The relationship between tumor angiogenesis, particularly microvessel density (MVD), and the histopathological subtypes of gastric adenocarcinoma remains underexplored. While extensively evaluated in malignancies such as breast cancer, the prognostic and therapeutic relevance of MVD in gastric cancer is yet to be fully established. Since angiogenesis plays a pivotal role in tumor growth, invasion, and metastasis, understanding its distribution across histological subtypes may guide the development of targeted anti-angiogenic therapies in gastric carcinoma.

This study aimed to assess the association between MVD, evaluated via PECAM-1 (CD31) immunohistochemistry, and the histological subtypes of gastric adenocarcinoma. The patients in our cohort were aged between 30 and 66 years, with a mean of 53.34 ± 8.38 years. The 50–59 year age group represented the largest proportion (54%) of cases, which is comparable to the findings of Pramanik et al., who reported a mean age of 55.3 years (SD ±12.71), with a range from 20 to 81 years in their Indian population [6]. In the present study, 64% were male and 36% were female, reflecting a male predominance consistent with Abdel-Aziz et al., who reported 62.5% male and 29.2% female among gastric carcinoma patients [14]. Similarly, Pramanik et al. observed a male-to-female ratio of 2.3:1 [7]. Histologically, 62% (31/50) of the cases were of the intestinal type, while 38% (19/50) were diffuse type, aligning with the results of Abdel-Aziz et al., who found 66.7% intestinal and 29.2% diffuse subtypes among their gastric adenocarcinoma cases [7]. Regarding angiogenesis, the mean MVD assessed through CD31 immunostaining was 17.44 ± 9.88, with a median of 13.50. This is in concordance with Badescu et al., who reported a mean MVD of 19.14 ± 4.25, with values ranging from 12 to 27 in 28 gastric tumor specimens [8]. In our series, high MVD was detected in 58.1% (18/31) of intestinal-type adenocarcinomas and in 36.8% (7/19) of diffuse-type tumors. Interestingly, this is somewhat divergent from Badescu et al., who reported higher MVD in diffuse-type gastric carcinoma compared to the intestinal type (33.4 vs. 26, p = 0.04) [8].

These findings underscore the potential difference of angiogenic activity across histological subtypes and highlight the utility of PECAM-1 based MVD assessment as a possible biomarker for tumor aggressiveness and treatment stratification in gastric adenocarcinoma.

## Conclusion

Based on the histopathological and immunohistochemical findings, it can be concluded that microvessel density (MVD) assessed by PECAM-1 (CD31) expression, is positively associated with the aggressiveness of gastric adenocarcinoma. A higher MVD indicates increased angiogenic activity, which contribute to more rapid tumor progression, particularly in intestinal subtypes. This study signifies the clinical relevance of tumor angiogenesis in gastric cancer and highlights the potential role of PECAM-1 based MVD assessment in classifying patients according to angiogenic burden. These findings may assist clinicians in identifying candidates for anti-angiogenic therapies, especially within the intestinal subtype of gastric adenocarcinoma, where targeted interventions aimed at disrupting tumor vascularization could prove beneficial in controlling disease progression.

## Notes

### Competing Interest Statement

The authors have declared no competing interest.

